# An ecophysiological model of plant-pest interactions: the role of nutrient and water availability

**DOI:** 10.1101/807941

**Authors:** Marta Zaffaroni, Nik J. Cunniffe, Daniele Bevacqua

## Abstract

Empirical studies have shown that particular irrigation/fertilization regimes can control pest populations in agroecosystems. This appears to promise that the ecological concept of bottom-up control can be applied to pest management. However, a conceptual framework is necessary to develop a mechanistic basis for empirical evidence. Here we couple a mechanistic plant growth model with a pest population model. We demonstrate its utility by applying it to the peach - green aphid system. Aphids are herbivores which feed on the plant phloem, deplete plants’ resources and (potentially) transmit viral diseases. The model reproduces system properties observed in field studies and shows under which conditions the diametrically-opposed plant vigour and plant stress hypotheses find support. We show that the effect of fertilization/irrigation on the pest population cannot be simply reduced as positive or negative. In fact, the magnitude and direction of any effect depends on the precise level of fertilization/irrigation and on the date of observation. We show that a new synthesis of experimental data can emerge by embedding a mechanistic plant growth model, widely studied in agronomy, in a consumer-resource modelling framework, widely studied in ecology. The future challenge is to use this insight to inform practical decision making by farmers and growers.

## Introduction

In agriculture, pest control mostly relies on the use of chemical pesticides. However, widespread application of agrochemicals carries an inherent environmental cost. There is also the significant challenge of declining efficacy due to the emergence and spread of insecticide resistance in pest populations (1). In recent decades, agroecology has developed as discipline which aims to provide alternatives to the use of chemicals in agronomy to control pest. The rationale is that ecological concepts and principles can be applied to control pest populations while reducing the use of chemicals (2). The concept of ‘bottom-up’ control, according to which population dynamics are driven by quantity and quality of resources, is particularly highlighted by agroecologists. There are a number of agricultural practices that can affect plant physiology and alter resources offered by plants to pests (3; 4). For example, fertilization modifies nutrient balance in plants, enhancing plant tissue nutritional status, and influences the synthesis of defence compounds (5). Similarly, irrigation controls plant vigour, phloem nutritional quality and viscosity, possibly regulating pest abundance (6; 7; 8; 9; 10).

Unfortunately, how pests might be affected by plant nutrient and irrigation status is far from obvious. Empirical evidence is ambiguous, potentially supporting diametrically-opposed hypotheses. On the one hand, the Plant Vigour Hypothesis (PVH) (11) argues that pest populations should increase most rapidly on vigorously growing plants (or organs), since these habitats provide more resources. In support of this hypothesis, there is some experimental evidence suggesting that practices such as fertilization and irrigation, or favourable conditions for plant growth such as increased organic soil fertility, can be associated with abundant pest populations (12; 13). On the other hand, the Plant Stress Hypothesis (PSH) (14) argues that pests perform better on stressed plants that would not have resources to deploy defences and/or whose nutritional quality might be enhanced. This as been determined experimentally to be the case for some aphid species feeding on plants subjected to controlled irrigation deficit (15; 16).

In order to efficiently use the concepts of bottom-up control in agroecology, it is necessary to shed light on the mechanisms that are responsible for the observed patterns. We require a unified conceptual framework sufficiently flexible for both the PVH and PSH hypotheses to find support. Developing and validating such a framework requires integration of information from field experiments with mathematical modelling. Experimental data is clearly necessary to test the validity of theoretical hypotheses, but is often extremely costly and time consuming to obtain. Mathematical modelling, particularly mechanistic models, represent a useful tool to investigate which processes can be responsible for the observed patterns and to explore the consequences of different agricultural practices (17).

Here we present a new, explicitly agro-ecological, model synthesising elements of models as commonly used within the separate disciplines of agronomy and plant pest dynamics. Agronomic models tend to empirically parametrize the detrimental effects of pests on plant biological rates (*e*.*g*. photosynthetic, growth, solutes transport). However such models invariably neglect the dynamical interaction between the plant (or some of its component parts) and the pest (see *e*.*g*. 18; 19; 20). That is, the impact of a pest on the plant is modelled by varying one or more plant parameters, according to the pest disturbance level with no further interaction or feedback. On the other hand, in ecology, there is a very broad literature of models on interactions (*e*.*g*. predation, consumption, competition etc.) between different species or organisms. These types of models have been widely used to study temporal and spatial dynamics in plant-pest (*e*.*g*. 21; 22; 23) and particularly plant-pathogen systems (*e*.*g*. 24; 25; 26). However these types of model usually present a simplistic description of the plant (but see (27)), which in turn limits the possibility to consider the effects of agronomic practices.

Aiming to bridge the gap between the classical agronomic and the ecological modelling approaches, here we couple a plant growth model, that describes carbon and nitrogen assimilation and allocation to shoot and root compartments of a plant, with a pest population model. With regard to the plant, we use the modelling framework proposed by Thornley in the early 70s (28), and refined in the following decades (29; 30; 31), which represents a cornerstone in plant and crop modelling. With regard to the pest, we propose a novel population model which includes intraspecific competition in which pest birth and mortality rates depend on resource availability and quality. Moreover, we assume that the presence of the pest can induce the plant to produce defensive traits or compounds (32). We demonstrate the utility of our model by applying it to the peach (*Prunus persica*) - green aphid (*Myzus persicae*) system. Aphids are specialized herbivores which feed on the phloem of vascular plants. This depletes plants’ resources, affecting growth and reproduction, as well as eventually impacting upon yield (33). Moreover, aphids are the most common vector of plant viral diseases and so can often cause indirect damage far exceeding direct impacts via herbivory (34). We use likelihood-based techniques to calibrate model parameters and select model assumptions against field data obtained under different conditions of irrigation and fertilization. The resulting model has the ability to reproduce different system properties observed in field studies, as well as showing under which conditions the PVH and PSH find more support. Our model also provides insights to conceive new targeted experiments to better understand this class of system and rethink the control of plant-aphid systems.

## Material and Methods

### Model outline and assumptions

The model, which describes the temporal variation, during a growing season, of plant dry mass (partitioned into shoots and roots, in turn composed of structural mass, carbon and nitrogen substrates), its induced defensive level and the aphid population dwelling on the plant, is schematically represented in Figure 1. According to Thornely et al’s seminal works (28; 17; 29; 30), carbon is assimilated from the atmosphere via photosynthesis and stored in shoots, as shoot carbon substrate (C_S_), or transported and then stored in roots as root carbon substrate (C_R_). Similarly, nitrogen is assimilated from the soil, stored in roots as root nitrogen substrate (N_R_), or transported and then stored in shoots as shoot nitrogen substrate (N_S_). Carbon and nitrogen substrates are utilized, in a fixed ratio, to constitute structural shoot (S) and root (R) dry mass. With respect to the original model of Thornley, we added the assumption that the constitution of new plant mass is regulated by changes in the photo-period (35). Such an assumption permits us to model the fact that perennial plants suspend growth, in favour of reserve constitution, before entering winter dormancy (36). The assimilation of substrate (*C_S_* or *N_R_*) per unit of plant organ (*S* or *R*) decreases with organ mass due to shoot self-shading and root competition for nitrogen and it is inhibited by substrate concentration in the organ (30).

**Figure 1:**
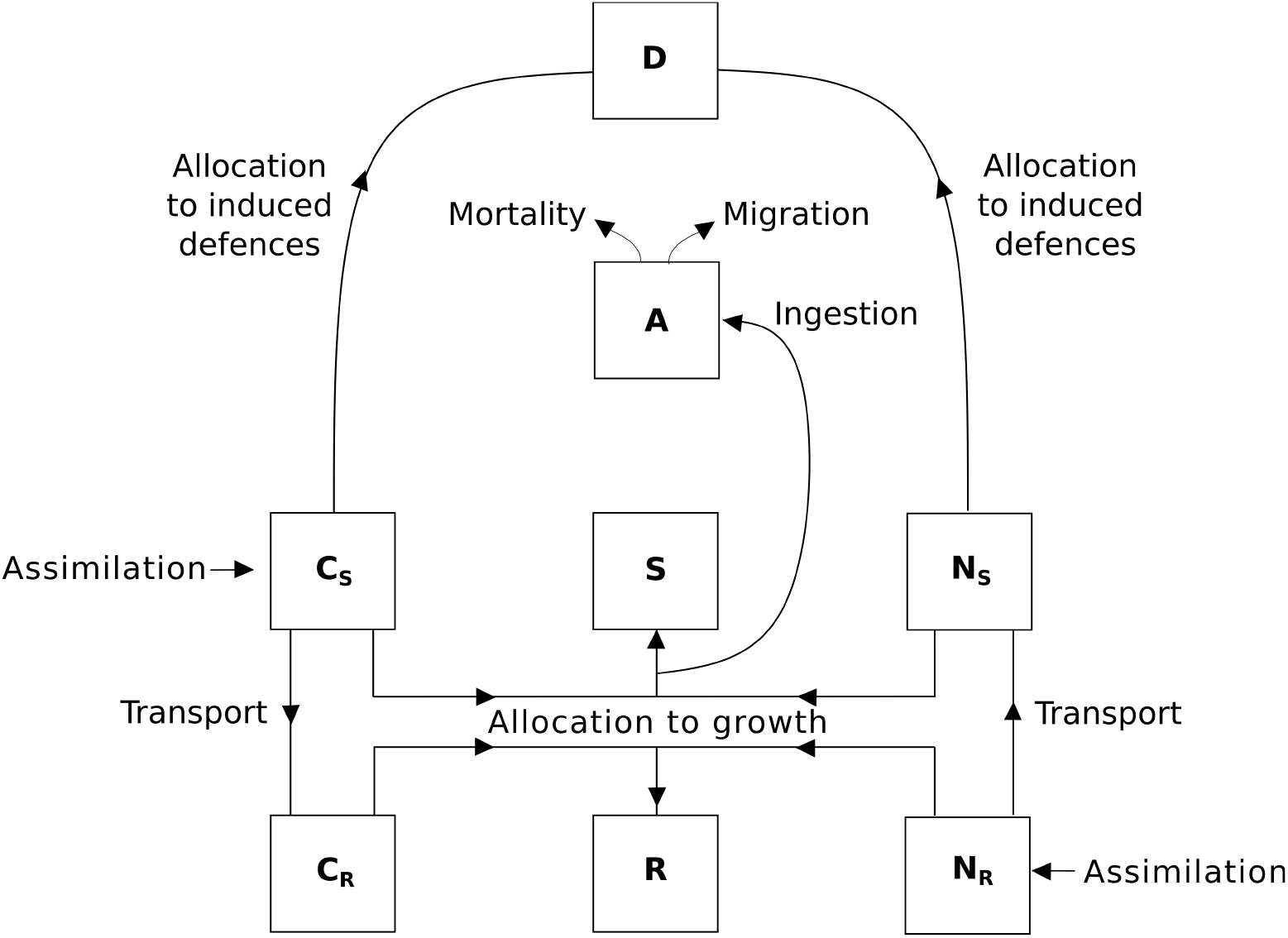
Schematic representation of the plant-aphid model where the plant is constituted by shoot (S) and root (R) structural dry mass, carbon (C_i_) and nitrogen (N_i_) substrates in shoots (i = S) and roots (i = R). The aphid population (A) intercepts a fraction of substrates allocated to constitute shoot structural mass and the plant diverts shoot substrates (carbon and nitrogen) to produce defensive compounds (D). More details are given in the main text.

We coupled the plant model of carbon and nitrogen assimilation and partitioning with an aphid population model by assuming that aphids, which penetrate growing shoots of the host plant with a stylet and feed on the phloem (37), intercept a fraction of the substrates (C_S_ and N_S_) directed towards the shoot structural mass compartment (S) to support their growth (38). We assume that aphids act in a scramble competition context (39) and therefore any aphid ingests its maximum daily amount of food when the per-capita available resource is sufficient, but that otherwise the resource is evenly shared among all the individuals. The aphid birth rate depends on the per-capita ingested food (40) and crowding can induce aphids to leave the plant (41).

We assumed that the infested plant can be induced to use carbon and nitrogen substrates to defend itself, to the detriment of growth (42; 37; 43). This can result in the production of chemical and/or in morphological and physiological changes that can reduce aphid accessibility to the resource (e.g. by phloem sealing) (44; 45) and/or decrease the rate at which ingested food is converted into progeny, e.g. by releasing toxic components in the sieve that can even repel or kill the aphid (32).

### Model equations

In quantitative terms, we describe the temporal variation of the plant-aphid system with the following system of ordinary differential equations.

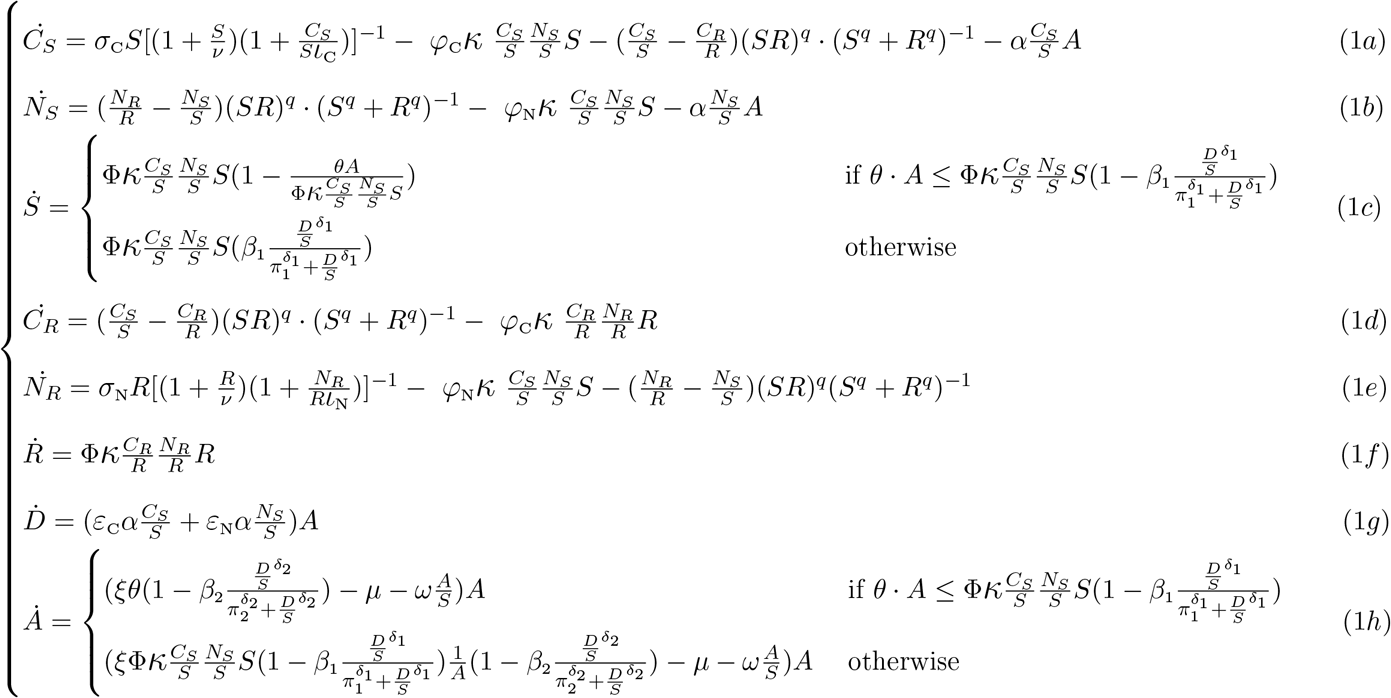

In our model *C*_*S*_, *N*_*S*_, *S*, *C*_*R*_, *N*_*R*_ and *R* are expressed in grams (g); *D* is expressed in an arbitrary defence unit (DU) and *A* in individuals (ind.); *t* represents the number of days (d) that have passed since the January 1^st^ of the year of the considered growing season. In equation 1a, 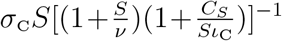 is the carbon substrate assimilated in shoots, 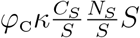 is the shoot carbon substrate allocated to shoot growth or reserves, 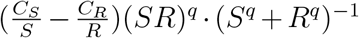 is the shoot carbon substrate transported toward roots and 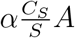 is the shoot carbon substrate diverted to defences, in a unit of time. In equation 1b, 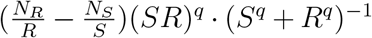 is the nitrogen substrate transported from roots towards shoots; 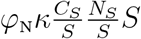 is the shoot nitrogen substrate allocated to shoot growth or reserves, and 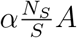 the shoot nitrogen substrate diverted to defences, with each of these quantities being measured as rates per unit of time. In equation 1c, the time dependent parameter 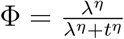 determines the suspension of plant growth driven changes in the photo-period The term 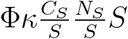 is the increase in structural shoot dry mass in the absence of any phloem withdrawal by the aphids, with *k* being the maximum rate of utilization of the substrates. The term 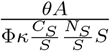 represents the fraction of substrates diverted from allocation to plant growth, because of ingestion by aphids, when aphid per-capita intake is limited by aphid maximum daily food intake, *θ*, and not by resource availability. The term 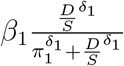 indicates the fraction of phloem that is protected by plant defences and therefore inaccessible for aphids. When aphid per-capita intake is limited by the resource availability, aphids ingest all the phloem they can access and the per-capita intake is reduced. The dynamics of the variables in the root compartments (*C*_*R*_, *N*_*R*_, *R*) follow similar rules as for assimilation of substrates, transport and allocation to root growth and we assumed that they are not directly affected by the presence of aphids. In equation 1h, we assume that the aphid birth rate is proportional to the per-capita food intake 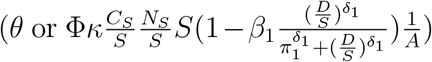 and that it can decrease due to a possible action of the defences. In other words, we assume that plant defences can determine an extra mortality rate, per unit of ingested food, modelled as 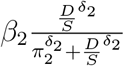. We modelled both the fraction of the phloem that can be protected and the phloem “toxicity” as an increasing function of the concentration of defences, 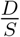. The shape of this function is given by the value of parameter *δ*_*i*_. Namely, if *δ_i_ >* 1 it is convex for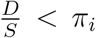 and concave for 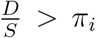, if 0 < *δ*_*i*_ < 1, it is strictly concave. The parameter *ω* is the strength of possible density dependent mechanisms inducing aphid migration. Details of the model variables and parameters are reported in Table 1.

**Table 1:**
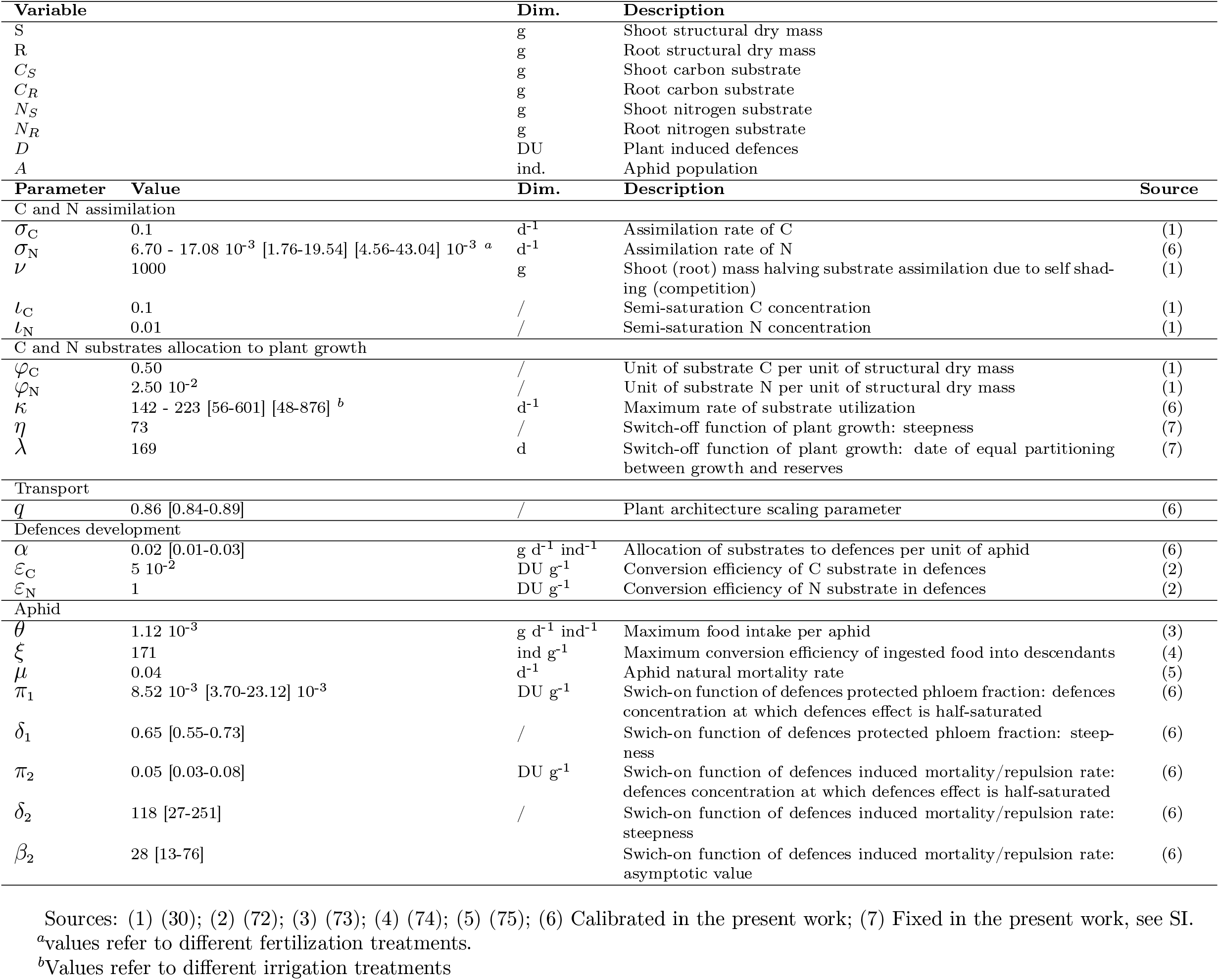
Model variables and parameters. For those parameters calibrated in the present work calibrated confidence interval of 99% are reported in brackets.

### Model calibration

We apply the model to a peach-green aphid system subjected to different fertilization and irrigation treatments. Details on the experimental data are reported in a previous work (8) and in the Supporting Information, SI. According to available data, we set initial conditions of the system at the first observation date (i.e. April 29^th^, 119^th^ day of the year 2013) (see SI). We set the value of model parameters according to information available from peer reviewed literature or experimental data whenever possible (Table 1 and SI). On the other hand, no information was available to *a priori* derive reliable estimates for parameters *σ*_N_ (net N assimilation rate) and *κ* (maximum rate of utilization of the substrates), which depend on environmental conditions that possibly varied in the different treatments; parameter *q*, affecting substrates transport within the plant and depending on the plant architecture (30), and six parameters relevant to the production of defences (*α*) and their effect (*π*_1_, *δ*_1_, *β*_2_, *π*_2_, *δ*_2_). We estimated these unknown parameters by minimizing a cost function expressed as the sum of two negative log-likelihood functions, computed with respect to observations of shoot dry mass and aphid abundance (see SI for details). We assessed the empirical distributions of calibrated parameters by making use of the moving block bootstrap (46). In particular, we reconstructed bootstrapped time series for each of the observed variables and we fitted the values of the unknown parameters. We repeated this process 1,000 times and we generated the 99% confidence intervals CI for each parameter via the percentile method (47).

### Model selection

To account for possible different mechanisms regarding aphid ecology, the plant response to aphid infestation and its consequences, we contrasted the ‘full’ model reported in eq.1 with a set of nested models lacking some processes (Fig. 2). Namely, the full model (M10) assumes that aphid crowding promotes aphid migration, that the plant produces defences that make a fraction of resources inaccessible to aphids and kill, or repel, aphid if ingested. Three models nested in M10 assume a crowding effect on aphid migration and the induced production of defences. Yet, they can differ regarding the effect of defences: killing/repulsion effect (M9), reduction of phloem accessibility (M8), or no effect (M7). There is also a simpler model neglect the production of defences (M6). We also considered five analogous models ignoring the effect of aphid crowding, *ω* = 0, (M1, M2, M3, M4, M5).

**Figure 2:**
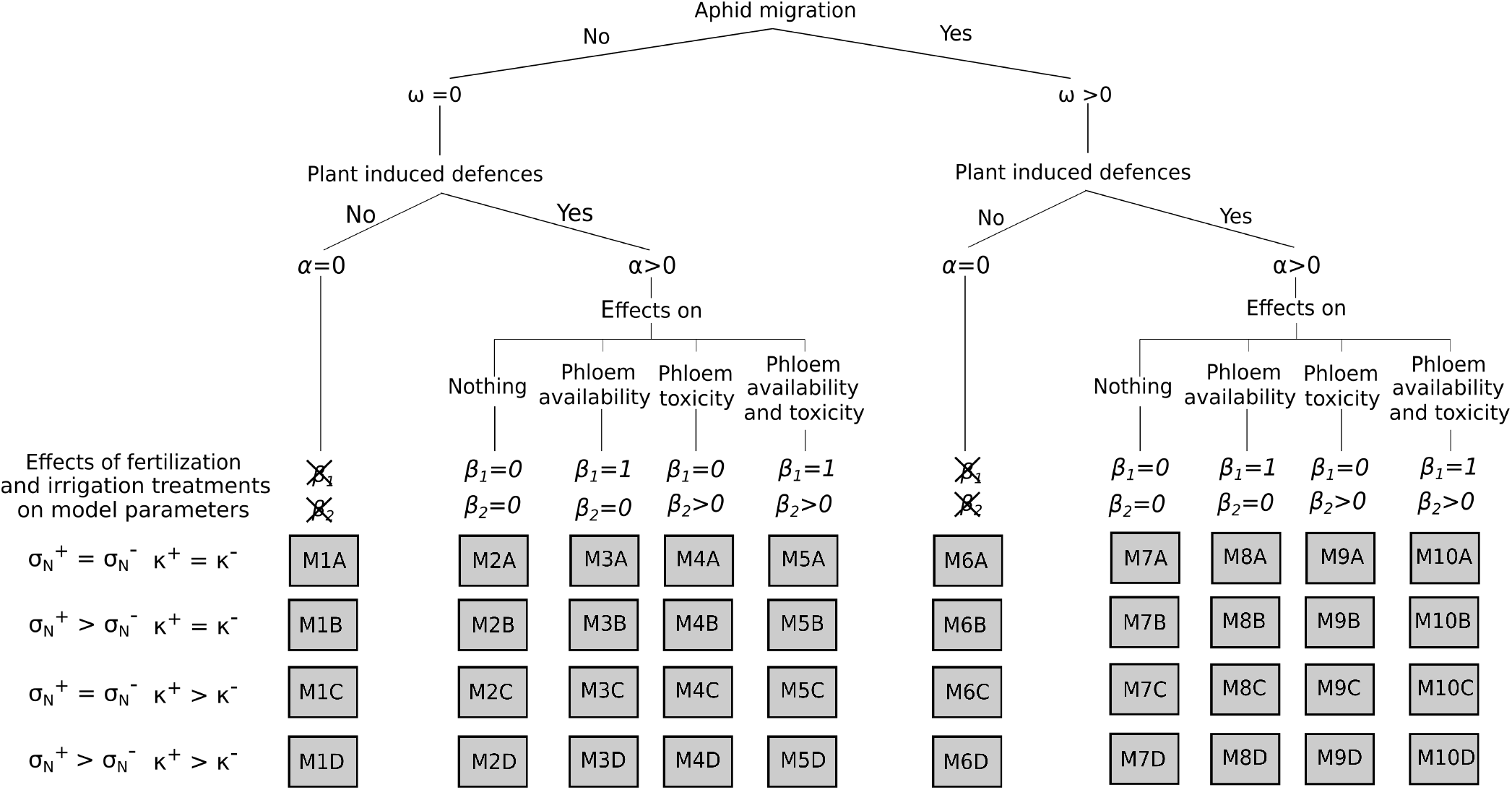
Schematic representation of the mechanisms considered in the different models *Mi* (*i ∈* [1, 10]) nested in eq.1: *i)* density dependent aphid migration (*ω*), *ii)* plant induced defences development (*α*) and *iii)* effect of induced defences on phloem availability to aphids (*β*_1_) and on phloem toxicity (*β*_2_). When the model parameter is set to zero, the relevant mechanism is ignored. Each model can be based on different hypotheses about the variation of the nitrogen assimilation rate *σ_N_* (equal (A, C) or different (B, D) across fertilization treatments) and the substrates utilization rate *k*(equal (A, B) or different (C, D) across irrigation treatments)

We tested if the effect of irrigation and fertilization can be represented in the models thorough a variation in some parameters *κ* and *σ*_N_. The rationale is that the rate of utilization of the substrates (parameter *k*) and the nitrogen assimilation rate (parameter *σ*_*N*_) are expected to decrease in water (48; 7) and nutrient (49; 50) stress conditions, respectively. We then contrasted each of the ten models assuming that *i) κ* and *σ*_N_ respectively vary with irrigation and fertilization treatments; *ii) κ* varies with irrigation and *σ*_N_ does not vary with fertilization; *iii) κ* does not vary with irrigation and *σ*_N_ varies with fertilization; *iv)* neither *σ*N nor *κ* vary with fertilization and irrigation. Therefore, we calibrated two values for nitrogen assimilation rate per unit of root 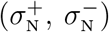 in cases *i* and *iii* and a unique value 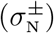 in cases *ii* and *iv*. Analogously, we calibrated two values for the allocation of substrates to plant growth (*κ*^+^ and *κ^−^*) in cases *i* and *ii* and a unique value (*κ^±^*) in cases *iii* and *iv*.

Overall, we compared 40 different models (Fig. 2), obtained by incorporating five hypotheses on plant defences, two on density dependence of aphid migration, and four on the effect of irrigation and fertilization, to one another. We selected the best model, that is the one providing the best compromise between goodness of fit to observed data and parsimony, through a model selection procedure based upon Akaike information criterion (51). See SI for details.

### Sensitivity Analysis

To assess the robustness of model outputs to uncertainty affecting model parameters, we performed a sensitivity analysis of the model to (small) perturbations of the default parameter values reported in Table 1. According to (17), we computed the sensitivity of the variable Y (where for Y we considered the maximum value of S, of A and of their ratio A/S over the growing season), to small variations of parameter *p_i_* as

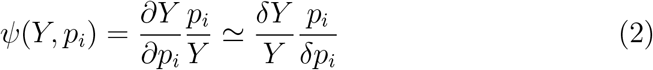

In practice, after having changed the value of each parameter by +5%, we computed the value of *ψ* and if *ψ*(*Y, p_i_*) *>* 1 we concluded that parameter has a more-than-linear effect on the variable.

### The role played by fertilization and irrigation

After having ascertained that parameters *κ* and *σ*_N_ are likely to vary with irrigation and fertilization treatments, respectively, we used the selected model to simulate the temporal dynamics of the system for different values of these parameters. This allowed us to perform an *in silico* experiment to explore whether or not the model was able to reproduce the observed empirical patterns that claimed support for the plant vigour or the plant stress hypotheses. The *in silico* experiment is intended to test if the aphid density is affected by the fertilization (or irrigation) treatment. We considered five levels for the fertilization treatment (i.e. *σ*_N_ equals to 0.0012, 0.0024, 0.012, 0.06 and 0.12 *d*^−1^) and five levels for the irrigation treatment (i.e. *κ* equals to 18, 36, 182, 910 and 1820 *d*^−1^) corresponding to very low - low - average - high - very high levels of fertilization (or irrigation). We varied the level of one treatment while keeping the other fixed at its average value thus obtaining nine different combinations of factorial levels. Since in real factorial experiments the number of replicates (i.e. different plant individuals) is limited, we chose to simulate ten replicates for each factors combination, which corresponds to a realistic experiment with 90 plants being monitored. We simulated ten possible trajectories of the system variables, for the same factors combination, by running the model with ten different parameter sets drawn from the empirical distribution obtained in the calibration process.

## Results

### Model calibration and selection

The best model (‘the model’, hereafter) assumes that *i*) aphid migration due to crowding can be neglected; *ii*) aphid presence induces the plant to divert resources from growth to defence; *iii*) defences reduce phloem accessibility to aphids and, at higher concentrations, make the phloem sufficiently toxic to kill or repel aphids (Fig. S1 in the SI); *iv*) the rates of nitrogen assimilation and substrates utilization differ for different levels of fertilisation and irrigation, respectively.

The model fitted all four data sets, reproducing the main observed temporal patterns and differences between treatments (Fig.3). Shoot growth is enhanced in *N* ^+^ treatments while the water treatment considered here plays only a relatively minor role. The time course for shoot mass is linear and followed by a stop towards the end of June. This is consistent with a potential exponential course, in the first part of the season (52), which has been prevented by the presence of the aphids. On the other hand, the stop in shoot growth at the end of June is induced by changes in the day-length. Note that parameter *ϕ*(*t*) = 0.5 for *t* = λ = 169, corresponding to June 18^th^. Aphid population growth is initially sigmoidal, followed by a decay towards the end of June when the plant growth is halted (Fig.3) and the concentration of defences attains the critical value of *π*_2_ = 0.03 − 0.08 which makes ingestion from the phloem detrimental rather than beneficial to aphids. The initial phase of aphid growth is enhanced in *N* ^+^ treatments characterized by more vigorous plants.

**Figure 3:**
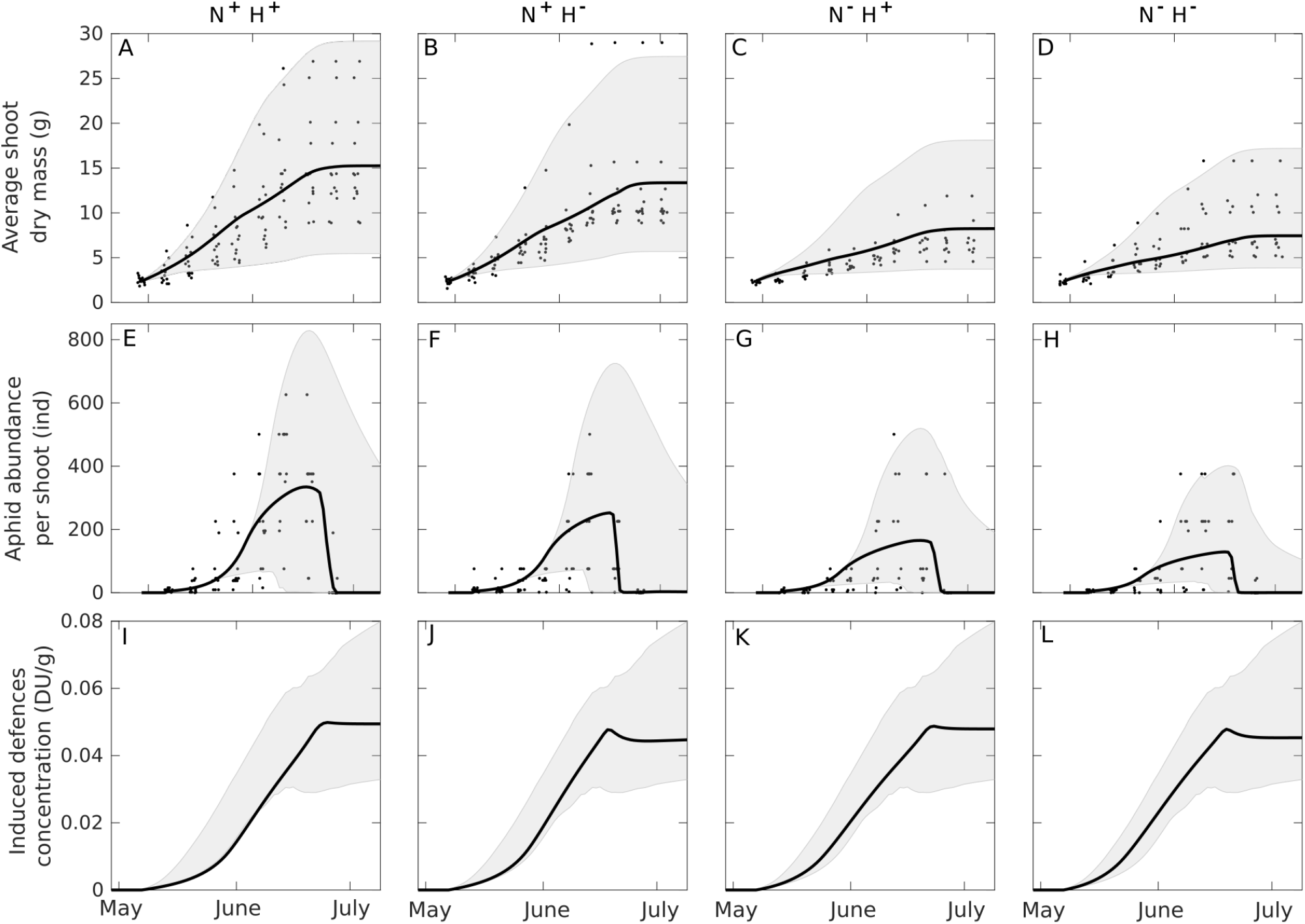
Observed (black points) and predicted (black lines) values of average shoot dry mass (top row), average aphid abundance per shoot (central row) and induced defences concentration (bottom row) under different fertilization and irrigation treatments: high fertilization and irrigation (A-E-I), high fertilization and low irrigation (B-F-J), low fertilization and high irrigation (C-G-K), low fertilization and irrigation (D-H-L). Grey shaded areas indicate the predicted 99% confidence bands.

The model gives biologically plausible parameter estimates (Table 1) and the estimated variability in parameters permits most of the variability observed in the data to be captured. The calibrated values of *σ_n_*, *k* and *q* are consistent with previously published values (i.e. *σ_n_* = 0.02*d*^−1^, *k* = 200*d*^−1^ and *q* = 0.67 − 1 in (30)). The estimated values of parameters *δ*_1_ < 1 and *δ*_2_ > 1 suggest that the fraction of phloem that is protected from aphid withdrawal quickly increases for low concentrations of defences, whereas the phloem toxicity is switched-on when the concentration of defences exceeds a threshold value (Fig. 1 in the SI). On the other hand, the model parameters relevant to the production of defences and their effect on aphids have no equivalent in the literature for a direct comparison.

### Sensitivity analysis

Ranked values of the sensitivity, *ψ*, of shoot production, and maximum aphid abundance and density to small changes in the parameter values are reported in Table S2 in the S.I. Negative values of *ψ* indicate a negative correlation between a change in a parameter value and the corresponding variable of interest. As expected, increasing the parameter λ results in an increase of shoot production, as it determines an increase of the growing season of 8.45d, being 0.05λ = 8.45, and consequently more resources to sustain a bigger aphid population, maintaining similar aphid densities. Similarly, an increase of *q* results in an increase of both shoot production and in the peak of aphid abundance and density, as it determines a more efficient transport of substrates C and N between roots and shoots. This translates into bigger plants able to sustain higher peaks of aphid population densities.

With the exception of *q* and λ, our sensitivity analysis indicates none of the model parameters has important (e.g. *ψ* > 1) consequences, indicating that the model is robust. However, our sensitivity analysis nevertheless provides some interesting insights. For instance, it shows that an increase in all those parameters positively related to the plant growth (*σ_c_, σ_n_, ι_C_, ι_N_, k, ν*) determine an increase in the maximum aphid abundance and, to a lower extent, in maximum aphid density. If the aphids were more efficient in converting food into progeny (higher *ξ*), aphid density would increase but the overall population abundance would diminish as the resource would be overexploited. An increase of the parameter *α*, determining a higher rate of resources devoted to defences, would have almost no effect on the shoot production but it would decrease aphid abundance and density. Yet, the plant could take advantage of a lower aphid abundance being aphids important vectors of viral diseases (37).

### The role played by fertilization and irrigation

Shoot growth follows a sigmoidal pattern and it increases with fertilization and irrigation (Fig. 4A-B). The concentration of carbon substrates in shoots varies between 3-23% during the growing season with peaks at its beginning, when plant growth is limited by the nitrogen supply, and at its end, when plant growth halts in response to day length decreases, but carbon assimilation continues. Carbon concentration is enhanced in stressful conditions (very low to low fertilization/irrigation treatments) that limit plant growth rather than carbon assimilation (Fig. 4C-D). The concentration of nitrogen substrates varies between 0.1-1.4 % during the growing season (Fig. 4E-F). It decreases in the first weeks of growth, but, in the case of very high/high fertilization, or very low irrigation, it increases until the second week of May. In fact, for high fertilization treatments, nitrogen is not initially consumed by plant growth which is limited by carbon supply and, for low watering, nitrogen concentration increases as plant growth is impaired while N assimilation is not. Peak concentration of defences is delayed in time for higher fertilization and irrigation (Fig. 4G-H). When plant is well watered, the time of the peak aphid population density is delayed by one week. This is due to the fact that defences need more time to reach significant concentrations in bigger plants (Fig. 4I-J). The positive effect of fertilization and irrigation upon aphid abundance becomes evident in the end of May. In the first part of the season, aphid density is enhanced by a low/average value of fertilization (or irrigation) while later in the season aphid density is higher in a well fertilized (irrigated) plant (Fig. 4K-L).

**Figure 4:**
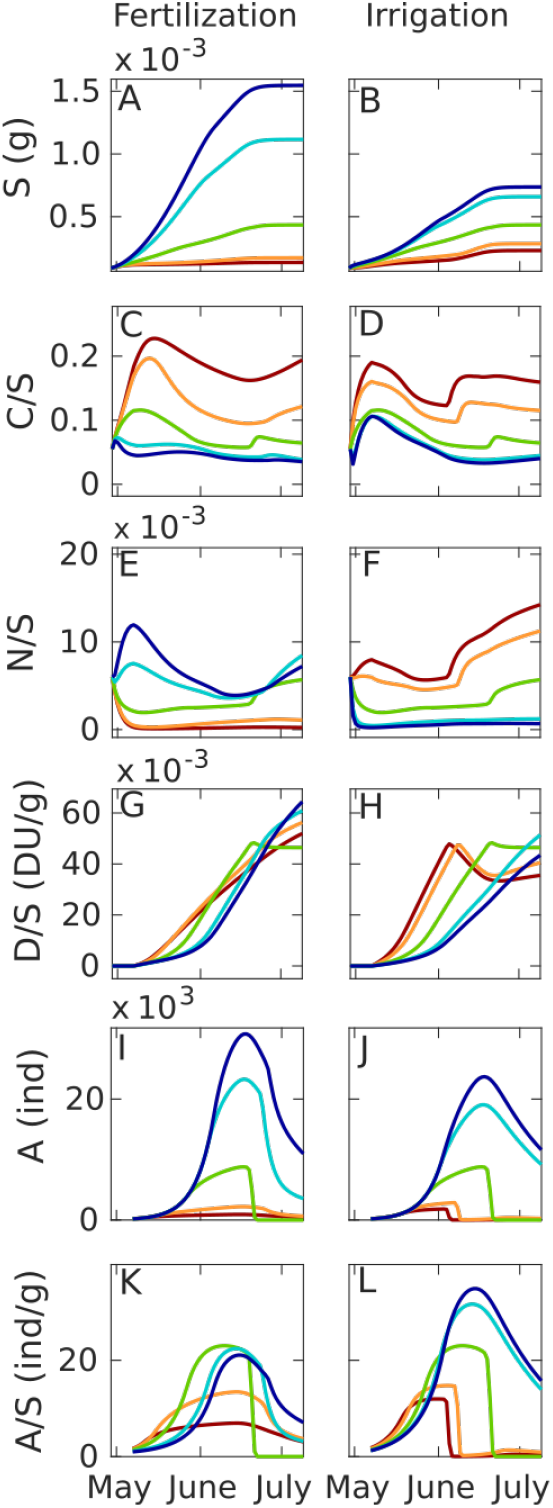
Simulated effect of fertilization (left column) and irrigation (right column) on the plant-aphid system: average shoot dry mass (A, B), carbon (C, D) and nitrogen (E, F) substrate concentration in shoots, defences concentration in shoot (G, H), aphid abundance (I, J) and density (K, L). Lines colour identifies fertilization (or irrigation) level: very low (red), low (orange), average (green), high (light blue), very high (blue).

The results of our virtual experiment show that one can draw very different conclusions depending on the considered fertilization/irrigation levels and the date of observations. For instance, one could infer that *i*) fertilization enhances aphid population by observing aphid density in the mid-late part of the season for very low to average values of fertilization (Fig. 5C-E); *ii*) decreases it, by observing aphid density in the early-mid season for average to very high values of fertilization (Fig. 5A-C); *iii)* has no effect, by observing aphid density early and late in the season, for high to very high values of fertilization (Fig. 5A-E). Similarly, different conclusions can be drawn regarding the effect of irrigation: positive (Fig. 5F), negative (Fig. 5B) or null (Fig. 5D, from average to very high values of irrigation).

**Figure 5:**
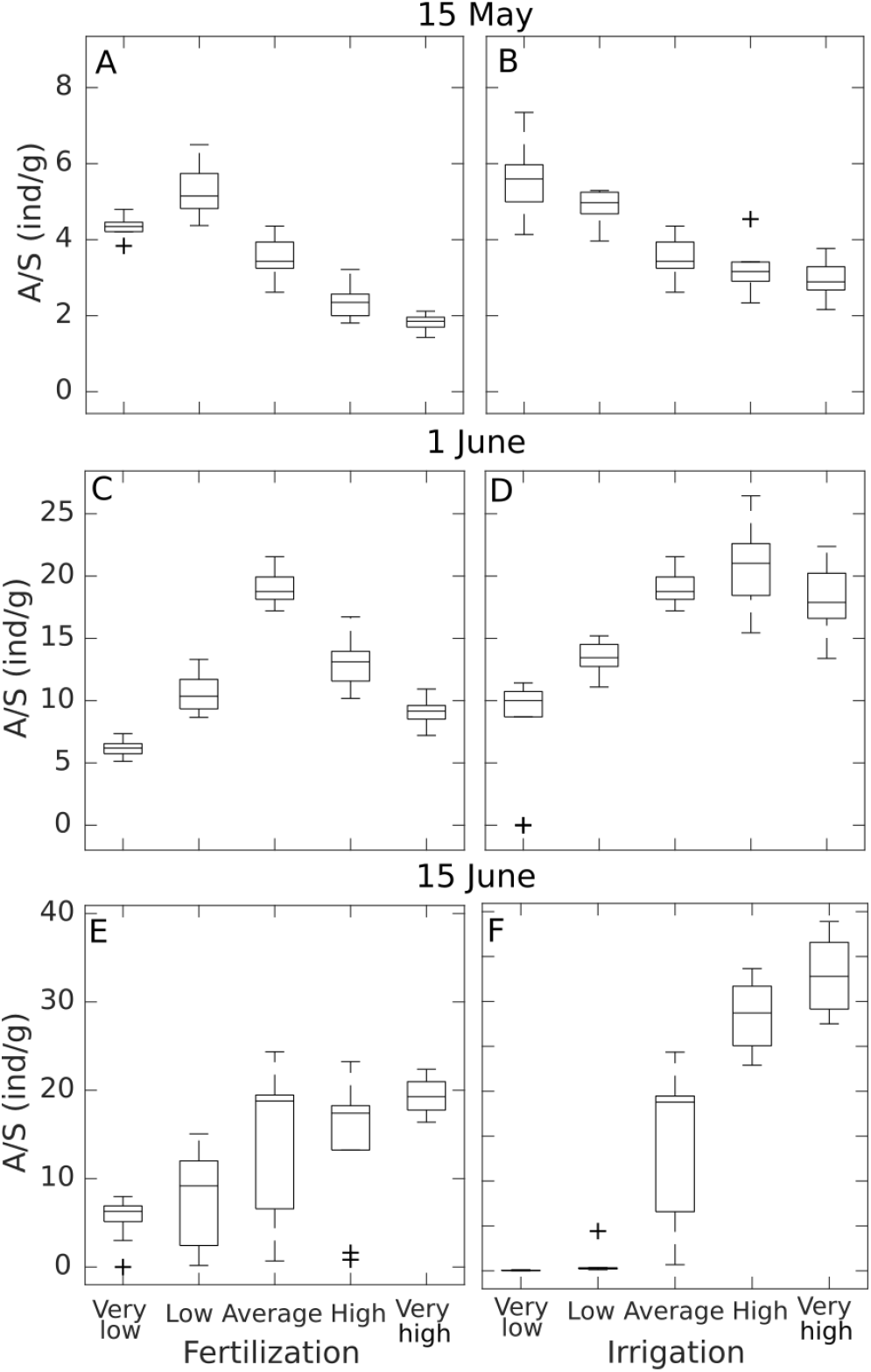
Simulated effect of fertilization (A, C, E) and irrigation (B, D, F) on aphids density on May 15^th^, June 1^st^ and June 15^th^. Boxes represent the first and third quartiles [25% and 75%] with a line inside indicating the median of ten simulated replicates of each treatment. The whiskers extend *±* 1.5 *×* the interquartile range (75^th^ percentile – 25^th^ percentile) from the third and first quartiles. Values outside the whiskers are considered outliers and plotted individually using the ‘+’ symbol.

## Discussion

In this work we showed that embedding a mechanistic plant growth model, widely studied in agronomy, in a consumer-resource modelling framework, widely studied in ecology, might be a promising approach for agroecology. We demonstrated the ability of such a novel approach in understanding the consequences of irrigation and fertilization treatments in a plant-aphid system.Yet, the proposed model has the ambition of being physiologically rigorous and general enough to be applied to different plant-pest systems and to incorporate the description of other agronomic practices.

### The selected model and model calibration and selection

A recent review (37) suggested that infested plants can put in place phloem-sealing mechanisms to interfere with aphids’ access to plant resources and produce a number of secondary metabolites (e.g. cardenolides, glucosinolates and benzoxazinoids) which, if ingested, impair aphid viability. Our study suggests that both defensive mechanisms are at play in the peach-green aphid system. According to our calibration, impairing phloem accessibility is the most effective at low defences concentration, while ‘intoxicating’ aphids is the most effective at higher concentration. This is in accordance with works on the arabidopsis-*Myzus persicae* system, for which reductions of aphids fecundity, up to 100%, have been reported in response to high concentrations of some plant defensive compounds (53; 54). The model application to a real study case subjected to different irrigation *×* fertilization treatments indicates that parameters relevant to plant nitrogen assimilation (*σ*_N_) and plant utilization of substrates (*κ*), originally proposed within a theoretical framework (28), can be linked to agronomic practices and then manipulated by the grower. However, in order to effectively use the proposed model to define effective agronomic recommendations, further studies on the response of the model parameters to effective practices are clearly required.

One of the main features of the peach-green aphid system is that, at the beginning of summer, aphid populations dwelling on peach trees drop. This occurs because aphids die, or abandon their primary host, or give birth to winged newborns that migrate to secondary herbaceous hosts (55). However, the underlying mechanisms triggering these processes are far from being clear. Our findings suggest that the reduction of resource availability, due to the investment in defensive traits and to photo-period driven interruption of shoot growth, along with the reduction of the phloem nutritional value, due to the accumulation of defensive compounds possibly toxic to the aphid, are the mechanisms most likely to be responsible for the observed patterns. In principle, the crash in aphid population could be due to other factors such as the arrival of predators attracted by high aphid density (56) or the possible reduction of the phloem nutritional value due to plant ageing (5). However, if the aphid population drop were driven by density dependent mechanisms, one would probably expect to observe fluctuations in the aphid population rather than a constant decline (57) and according to (58) there is no evidence that aphidophagous predators play an effective role in regulating aphid abundance. Moreover, in previous modelling works, it has been shown that observed population trends in different aphid species could be reproduced by considering a per capita death rate positively related to the aphid cumulative population size (59; 60; 61). Such a relationship coherently emerges as a property of our model if the pest presence induces the plant to produce defences that accumulate, and not if the phloem nutritional value declines throughout the season, independently from aphid presence.

Performing experiments to find correct numerical values for parameters of biological models is virtually impossible because many parameters cannot be directly measured. For this reason, we were forced to numerically calibrate nine parameters via our likelihood-based model fitting procedure. However, biologically plausible parameter estimates and good fitting does not guarantee that parameter estimates are correct, due to possible correlations among the parameters (62) and model identifiability problems that can arise due to an imbalance between model complexity and available data (63). The proposed modelling framework would therefore enormously benefit from experimental works dedicated to the measurement, or at least a sound assessment, of some model parameters. Despite the importance of the parameter *q* in Thornley’s models, we found no studies on its assessment. Similarly, although it is well known that a plant can divert resources from growth to defence (64), we found no quantitative relationships relevant to the cost of making defences (parameter *α* in our model) in terms of growth loss, neither between the presence of defences and pest performances. Our exercise provides a preliminary assessment of these parameters that need to be confirmed or confuted by dedicated field and/or laboratory works.

### The role played by fertilization and irrigation

Variations in plant growth, and in the concentration of C and N substrates in plant tissues, for different levels of fertilization and/or irrigation are well acknowledged (65; 48) and they have already been shown to be emerging properties of the original model for plant growth used in this work (28). Our pest-plant model maintains these properties (Fig. 4A-B-C-D-E-F) and allows further insights regarding the variations observed in aphid population. The aphid population response to fertilization and irrigation has been explored in a number of empirical works not providing a straightforward picture. Some authors observed no effect of fertilization in the wheat-Russian wheat aphid system (66), or negative effects of irrigation in the apple-rosy apple aphid and in the cotton-cotton aphid systems, respectively (67; 68). Other authors observed the highest aphid abundance at an average level of fertilization, and no effect of irrigation, in the chrysanthemum-cotton aphid system (69). The intrinsic rate of oat aphid population increase in three grass species was observed to be favoured by irrigation (70). On the other hand, aphid population was observed to be maximal for moderate water stress in the cabbage-green aphid and cabbage-cabbage aphid systems (16), and in one out of three genotypes tested for the poplar-wolly poplar aphid system (71). Our model, parametrized for the peach-green aphid system, shows that all these apparently contrasting empirical evidences can emerge from the same biological principles governing plant-pest dynamics and that both plant vigour and plant stress hypotheses can find support when observing a plant-pest system evolving in time and subject to different level of changes in the environment conditions. The aphid population dynamics reproduced by our model (Fig. 5) indicate that the effect of fertilization and irrigation on the pest population cannot be simply reduced as positive or negative In fact, its sign and strength depends on the considered levels of fertilization/irrigation and on the date of observation along the growing season. The contribution of our work is to show how a new synthesis of the experimental data can emerge by using mechanistic modelling. The challenge for our future work is to show how this insight – as well as the model developed here – can be used to inform practical decision making by farmers and growers.

## Aknowledgments

The field work to create the dataset used in this work was funded by the ARIMNET (ANR-12-AGR-0001) ‘APMed’ project (Apple and Peach in-Mediterranean orchards) and the ‘RegPuc’ project (Quelles stratégies d’irrigation et de fertilisation pour réguler les populations de pucerons verts en vergers de pêcher). The PhD grant of MZ is funded by the PACA region (Provence-Alpes-Côtes d’Azur) and INRA.

