## Supplementary Information for "An ecophysiological model of plant-pest interactions: the role of nutrient and water availability"

### Electronic Supplementary Material

#### Available data

Data come from 44 peach *Prunus persica* trees cultivated at the Institut National Researche Agronomique (INRA) station of Avignon (southern France, 43°60' N, 4°49' E). The plants were grafted in February 2011 and the experiment took place in spring 2013, from the end of April (corresponding to approximately 30 days after bloom) to the beginning of July (corresponding to the end of vegetative growth). Plants were subjected to four different treatments obtained by combining two levels of fertilization ($N^+$ and $N^-$) and two levels of irrigation ($H^+$ and $H^-$). On May 2$^{\text{nd}}$, two growing shoots per plant were selected and infested with five wingless adult aphid females each. The shoot growth and the aphid infestation level was weekly measured on each plant. Shoot growth was measured by counting the number of leaves per shoot. Aphid infestation was measured by assigning to each shoot an infestation class: C0 (no aphids), C1 (1-5 ind.), C2 (6-25 ind.), C3 (26-125 ind.), C4 (126-625 ind.) and C5 ($>$ 625 ind.). Available data are reported in files *shoot_ data.txt* and *aphid_ data.txt*. Details on the experiment are reported in the original paper of (1).

We computed dry shoot $S(t)$ mass as a function of the counted shoot leaves $n_S(t)$ via the allometric relationships $S(t) = 0.26 \cdot n_S(t)$ where $S$ is expressed in grams (2; 3). We computed aphid abundance $A(t)$ from the measured infestation class by setting it equal to 3, 16, 76, 376 and 625, respectively for classes from C1 to C5. We scaled up all the measures at the shoot level to the whole plant level by multiplying them by the number of growing shoots per plant.

#### Model initialization and off-line calibration

In order to run the model (i.e. integrate the system of ODE's) one must define the initial time $t_0$ and the corresponding values of the model state variables. We set $t_0$ to April $29^{th}$ (i.e. $119^{th}$ day of year DOY) and $S(t_0) = 19.7$ g and $A(t_0 + 3) = 41.5$ ind., per plant, according to field observations. Following (4), we assumed that at the beginning of the vegetative season, root:shoot ratio equals equals 0.3 and that the concentration of substrates in shoots and roots is 5.5% for C and 0.6% for N. We then set $R(t_0) = 5.9$ g, $C_S(t_0) = 1.09$ g, $N_S(t_0) = 0.12$ g, $C_R(t_0) = 0.33$ g, $N_R(t_0) = 0.04$ g. We set $D(t_0) = 0$ by assuming that induced defences are not present before infestation.

We used available data to assess the time at which the plant stopped growing. For each pairs of consecutive observation dates we evaluated via a t-test the probability $P$ that the means of the shoot dry masses were equal. If the plant has significantly grown in the considered period, the $P$ level will be $< 0.01$. We assumed that the first date of the pair for which $P > 0.01$ (i.e. $t$=169) corresponds to the parameter $\lambda$ of the switch off function $\Phi = \frac{\lambda^\eta}{\lambda^\eta + t^\eta}$. We assumed that at the first date $t^*$ of the pair for which $P > 0.05$ (i.e. $t*$=176) the value of $\Phi$ drops to a critical value $\Phi^* = 0.05$. After some algebra, one obtains $\eta = \frac{ln(\frac{1}{\phi^*} - 1)}{ln(t^*) - ln(\lambda)} = 73$.

#### Model calibration and selection

We estimated model unknown parameters by minimizing a cost function $C$ expressed as the sum of two negative log-likelihood functions $C = -(\ell_y + \ell_x)$ with:

$$\ell_i = -N_i log(\sqrt{2\pi\sigma_i{}^2}) - \frac{1}{2\sigma_i{}^2} \sum_P \sum_t (i_{P,t} - \hat{i}_{P,t})^2$$

where $i_{P,t}$ are the values of the average shoot dry mass ($i = y$) and the average aphid abundance per shoot ($i = x$) observed on the plant $P$ at time $t$, total samples size equal to $N_i$ and $\hat{i}_{P,t}$ are the corresponding values simulated by the model.

We assumed that the errors between each observation and the corresponding value estimated by the model follow a Gaussian distribution with mean 0 and unknown variance $\sigma_i{}^2$. To derive the log-likelihood functions, we assumed that the error structure is additive. We found the set of parameters that minimized the cost function $C$ using the Matlab command "fminsearch" (Nelder-Mead algorithm).

Once we calibrated the 40 possible models (see Fig.2), we selected the best as the one leading to the best compromise between goodness of fit and parsimony in the number of calibrated parameters. According to the Akaike information criterion (5), for each model we computed a value of $AIC = 2C + 2n_p$, where $n_p$ is the number of calibrated parameters. We ranked the models according to their $AIC$ value (Table S1). We computed the $AIC$ differences ($\Delta AIC_i$) between the $AIC$ value of the $i^{th}$ model and the minimum AIC among all considered models. According to (6), models with $\Delta AIC_i < 2$ can be considered as equivalent(7; 8) and, among equivalent models, we selected the simplest one (i.e. the one with less estimated parameters) as the best.

### Figures

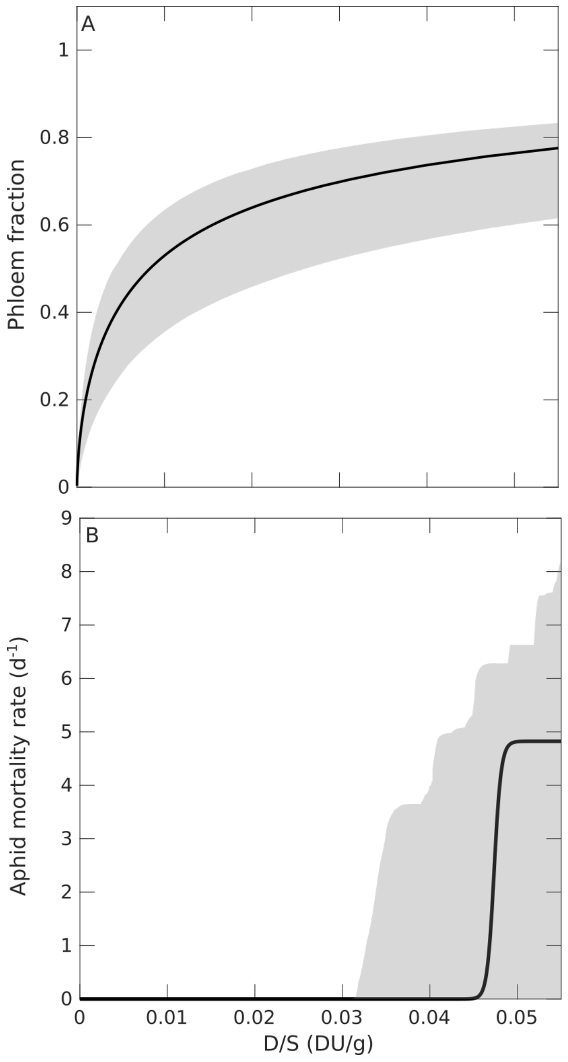


Figure S1: Effect of the concentration of plant defences on (A) Fraction of phloem that is unavailable to aphids and (B) aphid mortality induced per mg of ingested phloem. Grey curves indicate 99% confidence bands due to the uncertainty in the estimate of parameters $\pi_1$ and $\delta_1$ (A), and $\pi_2$, $\delta_2$ and $\beta_2$ (B).

### Table

Table S1: Comparison among candidate models for the plant-aphid system. For any model it is indicated with its identifier ID (see main text and in Fig. 2 for details), its complexity assessed by the number of calibrated parameters $n_p$; its Akaike score $AIC$; its $\Delta AIC_i$ computed as the difference between its $AIC$ and the lowest obtained from all the models i.e. $AIC = 6519.0$.

| ID | $n_p$ | $AIC$ | $\Delta AIC_i$ | ID | $n_p$ | $AIC$ | $\Delta AIC_i$ |
|---|---|---|---|---|---|---|---|
| M10D | 12 | 6519.0 | 0.0 | M8A | 7 | 6751.6 | 232.6 |
| M5D | 11 | 6520.8 | 1.8 | M2D | 6 | 6756.2 | 237.2 |
| M8D | 9 | 6570.8 | 51.8 | M4D | 9 | 6762.2 | 243.2 |
| M10B | 11 | 6576.1 | 57.1 | M3C | 7 | 6773.4 | 254.4 |
| M5B | 10 | 6590.5 | 71.5 | M2B | 5 | 6775.4 | 256.4 |
| M3D | 8 | 6624.5 | 105.5 | M4B | 8 | 6781.3 | 262.3 |
| M7D | 7 | 6628.4 | 109.4 | M4C | 8 | 6785.0 | 266.0 |
| M6D | 6 | 6632.1 | 113.1 | M3A | 6 | 6786.7 | 267.7 |
| M9D | 10 | 6634.2 | 115.2 | M6C | 5 | 6794.0 | 275.0 |
| M3B | 7 | 6641.5 | 122.5 | M7C | 6 | 6795.5 | 276.5 |
| M8B | 8 | 6641.9 | 122.9 | M6A | 4 | 6798.5 | 279.5 |
| M7B | 6 | 6646.4 | 127.4 | M7A | 5 | 6800.5 | 281.5 |
| M9B | 9 | 6651.6 | 132.6 | M2C | 5 | 6865.2 | 346.2 |
| M6B | 5 | 6696.0 | 177.0 | M2A | 4 | 6871.9 | 352.9 |
| M9C | 9 | 6708.6 | 189.6 | M4A | 7 | 6877.0 | 358.0 |
| M10C | 11 | 6712.6 | 193.6 | M9A | 8 | 6878.7 | 359.7 |
| M8C | 8 | 6721.6 | 202.6 | M1B | 4 | 7216.0 | 697.0 |
| M5C | 10 | 6727.9 | 208.9 | M1D | 5 | 7228.4 | 709.4 |
| M10A | 10 | 6742.9 | 223.9 | M1A | 3 | 7241.7 | 722.7 |
| M5A | 9 | 6746.8 | 227.8 | M1C | 4 | 7262.4 | 743.4 |

Table S2: Numerical approximation of the sensitivity, $\psi$ (see eq.2 in the main text), of the maximum value of shoot mass (S), aphid abundance (A) and density (A/S) to small changes in the parameters.

| | $\psi(S)$ | $\psi(A)$ | $\psi(A/S)$ |
|---|---|---|---|
| $\delta_2$ | 0.00 | 0.00 | 0.00 |
| $\delta_1$ | 0.01 | -0.35 | -0.27 |
| $\beta_2$ | 0.02 | 0.00 | 0.00 |
| $\mu$ | 0.02 | -0.31 | -0.32 |
| $\alpha$ | 0.02 | -0.65 | -0.43 |
| $\pi_1$ | -0.05 | 0.07 | 0.26 |
| $\eta$ | -0.06 | -0.01 | 0.01 |
| $\epsilon_N$ | 0.15 | -0.10 | -0.10 |
| $\nu$ | 0.17 | 0.10 | 0.05 |
| $\xi$ | -0.18 | -0.17 | 0.72 |
| $\kappa$ | 0.22 | 0.46 | 0.23 |
| $\epsilon_C$ | 0.25 | -0.12 | -0.16 |
| $\pi_2$ | -0.25 | 0.00 | 0.00 |
| $\theta$ | -0.25 | -0.68 | 0.19 |
| $\iota_C$ | 0.26 | 0.21 | 0.06 |
| $\iota_N$ | 0.29 | 0.27 | 0.04 |
| $\varphi_C$ | -0.53 | -0.71 | -0.15 |
| $\sigma_N$ | 0.63 | 0.64 | 0.06 |
| $\sigma_C$ | 0.65 | 0.63 | 0.16 |
| $\varphi_N$ | -0.66 | -0.76 | 0.03 |
| $q$ | 1.56 | 1.79 | 0.33 |
| $\lambda$ | 5.87 | 1.31 | 0.04 |
